# Rapid assessment of nitrification inhibitor efficacy, competitiveness, and specificity using microrespirometry

**DOI:** 10.64898/2026.06.25.734519

**Authors:** Christopher J. Sedlacek, Katharina Klawatsch, Bianca Lang, Elosie Atkinson, Frida Brandner, Ayla Horuz, Adrienn Markesz, Lucia Fuchslueger, Andrew T. Giguere, Petra Pjevac

## Abstract

Nitrification inhibitors are applied to reduce nitrogen losses and greenhouse gas emissions from fertilized agricultural ecosystems. However, their characterization is typically focused on determining effective inhibitor concentrations from growth or substrate conversion assays that are time-intensive and provide limited mechanistic resolution. Here, we present a microrespirometry (MR)-based workflow for rapid mechanistic characterization of nitrification inhibitors using oxygen consumption as a real-time readout for metabolic activity. The workflow enables the simultaneous assessment of inhibitor efficacy, competitiveness, and enzyme specificity within a single experimental setup, as sequential substrate and inhibitor additions enable direct discrimination between competitive and non-competitive inhibition and between ammonia monooxygenase-specific and broader respiratory inhibition. As a proof of concept, we evaluated three known nitrification inhibitors phenylacetylene (PA), nitrapyrin (NP), and dicyandiamide (DCD) using the ammonia-oxidizing bacteria *Nitrosomonas europaea* and *Nitrosospira multiformis*, the complete ammonia oxidizer *Nitrospira inopinata*, and the nitrite oxidizer *Nitrospira moscoviensis*. We also compared the results from the MR-based inhibition workflow with those from a conventional growth-based approach and observed a poor correlation between results for inhibitors that are not fully enzyme specific. In conclusion, this work establishes MR as a rapid and versatile platform for the mechanistic screening of novel potential nitrification inhibitors. MR assays reproduce known inhibitory responses while substantially reducing experimental time and increasing mechanistic resolution compared to other assays types. Additionally, we provide the first pure-culture characterization of PA, NP, and DCD efficacy and inhibition mechanisms in a complete ammonia oxidizer, *N. inopinata*.

## Introduction

Nitrification is the microbially mediated conversion of ammonia (NH_3_) to nitrite (NO_2_^-^) and nitrate (NO_3_^-^), and is one of the key processes in the global nitrogen (N) cycle. It is predominantly performed by aerobic ammonia-oxidizing bacteria (AOB), archaea (AOA), nitrite-oxidizing bacteria (NOB), and the more recently discovered complete ammonia oxidizers (comammox) (Treusch et al., 2005; Könneke et al., 2005; Daims et al., 2015; van Kessel et al., 2015; Daims et al., 2016; Kuypers et al., 2018). Industrial nitrogen fixation via the Haber-Bosch process has caused massive changes to the global nitrogen cycle, increasing the flow of nitrogen through the nitrification pathway (Rockström et al., 2009; Fowler et al., 2013). Most industrially synthesized NH_3_ is used to produce ammonium (NH_4_^+^)-based fertilizers, which accelerate nitrification rates in agricultural soils (Beekman et al., 2018). The produced NO_3_^-^ is more readily leached from soils than NH_4_^+^ and is a driver of eutrophication - the nutrient-induced increase of primary production in aquatic ecosystems that has devastating consequences for ecosystem health (Romanelli et al., 2020). In addition, nitrification and the subsequent microbial reduction of NO_2_^-^/NO_3_^-^ lead to increased NO and N_2_O emissions, which contribute to ozone depletion and atmospheric warming (Hu et al., 2015). Therefore, interventions to suppress nitrification in fertilized agricultural ecosystems have become a widely studied management practice in an attempt to regulate and mitigate the environmental impacts of nitrogen fertilization (Gioacchini et al., 2002; Lam et al., 2017; Coskun et al., 2017; Beekman et al., 2024).

To combat nitrogen losses and improve crop nitrogen use efficiency (NUE), novel technological and strategic advances in fertilizer application and new agricultural practices are continuously being developed (e.g. Xiang et al., 2008; Zebarth et al., 2009; Wu et al., 2021). The use of compounds that suppress ammonia oxidation (i.e., nitrification inhibitors), whether applied separately or combined with nitrogen-based fertilizer in the form of coated, slow-release fertilizer granules is one of the most promising ways to regulate nitrogen cycling in agricultural ecosystems (Subbarao et al., 2006a; Linquist et al., 2013; Dimkpa et al., 2020). However, to maximize efficient nitrification inhibitor use, their specificity, efficacy, and mode of inhibition must be elucidated, along with any off-target effects on the metabolic activity of other soil-dwelling microorganisms. This mechanistic characterization is necessary to ensure that nitrification inhibitors work efficiently but do not pose a risk to the broader soil health or ecosystem functioning (e.g., Gao et al., 2021; Woodward et al., 2021).

An ideal nitrification inhibitor inhibits ammonia oxidation non-competitively, irreversibly, with high efficacy, and in an enzyme-specific manner. To date, mechanistic studies of nitrification inhibitors have been primarily performed by growth and substrate conversion assays with pure cultures of nitrifying microorganisms (e.g., Subbarao et al., 2006b; Lehtovirta-Morley et al, 2013; Shen et al., 2013; Papadopoulou et al., 2020; Wright et al., 2020; Beeckman et al., 2022; Otakka et al., 2022) or through substrate conversion assays with soil communities (e.g., Keeney, 1980; Shi et al., 2017; Ma et al., 2021, Rojas-Pinzon et al., 2024). Growth, or lack thereof, provides insight into whole-cell inhibition and cytotoxicity, but such experiments with slow-growing nitrifying microorganisms are time and resource-intensive. In addition, growth inhibition alone provides little information about the competitiveness, reversibility or specificity of inhibitors. Substrate conversion assays, on the other hand, can be performed independent of growth, in a shorter time frame, and in a manner that allows more direct evaluation of the mode of inhibition (e.g., Wright et al., 2020; Taylor and Mellbye, 2022). Subsampling of such experiments allows for the generation of time-resolved substrate conversion datasets in the presence of inhibitors. However, neither growth nor substrate conversion assays provide real-time continuous measurements, nor do they offer a simple opportunity to determine efficacy, competitiveness, and enzyme specificity in parallel.

An alternative approach that directly addresses the shortcomings of growth and substrate conversion assays is to monitor oxygen (O_2_) consumption during substrate conversion. Measurement of O_2_ consumption (i.e., respirometry) is an almost universally applicable, sensitive, non-destructive, continuous, real-time measurement used to monitor the rate of metabolic activity within aerobic organisms in the absence of appreciable growth (e.g., Scholander et al., 1952; Owen and Legan, 1987; Spanjers et al., 1996; Young and Cowan; 2004). Respirometry is a well-established microbiology technique for cytotoxicity testing and metabolic assays (e.g. Chandran et al., 2008; Esquivel-Rios et al., 2014), although it has not been widely used to study the response of nitrifiers and other microorganisms to nitrification inhibitors (Voysey and Wood, 1987; Lontoh et al., 2008; Dalkidis et al., 2026). Respirometry can be performed in sealed, small volumes of microbial cultures using miniaturized electrochemical (Clark-type) oxygen microsensors or microoptodes (Revsbach, 1989; Klimant et al., 1997; Kühl et al., 2005), and in such setups is referred to as microrespirometry (MR). Importantly, MR allows for parameters other than oxygen concentration, such as temperature, pH, substrate, product, and inhibitor concentrations to be considered in a single experimental setup (e.g., Jung et al., 2022). Additionally, MR is a continuous real time assay format which eliminates the need to assume linear reaction rates between time points (Tonge, 2019). Thus, MR provides a highly efficient methods platform for the screening and mechanistic characterization of novel potential nitrification inhibitors.

Here, we present a straightforward MR workflow to assess the efficacy, competitiveness, and specificity (mode of action) of nitrification inhibitors in a single experiment. As proof-of-concept, we evaluated three known synthetic nitrification inhibitors: phenylacetylene (PA), nitrapyrin (NP), and dicyanamide (DCD) (e.g. Bedard and Knowles, 1989 and references therein; Vannelli et al., 1996; Lontoh et al., 2000; Wright et al., 2020). We demonstrate how MR is used to determine the inhibitory concentration (i.e. efficacy) for each inhibitor for several pure cultures of aerobic ammonia-oxidizing microorganisms: *Nitrosomonas europaea* Nm50 (AOB), *Nitrosospira multiformis* NI2 (AOB) and *Nitrospira inopinata* (comammox). Notably, this is the first published characterization of commercially used nitrification inhibitors on a pure comammox isolate, *N. inopinata* (Kits et al., 2017). In analogous MR assays with the NOB *Nitrospira moscoviensis* we demonstrate the use of MR to evaluate possible general cytotoxic effects of the inhibitors. Highlighting the versatility of the MR assay, we determined inhibitor competitiveness and inhibitor specificity for the ammonia monooxygenase (AMO) enzyme, which catalyzes the first enzymatic step of nitrification. Here, fully inhibited cultures were supplemented with additional substrate, both ammonium and subsequently hydrazine (N_2_H_4_). Hydrazine is a substrate analog for the hydroxylamine oxidoreductase (HAO) enzyme that catalyzes the second, AMO-independent enzymatic reaction in the ammonia oxidation pathway (Böttcher and Koops,1994; Vajrala et al., 2013). The resulting respiratory responses, or lack thereof, reveal inhibitor competitiveness and specificity. Finally, MR assay-derived efficacy results were compared with those obtained through conventional batch-culture growth assays with the same ammonia-oxidizing cultures, to enable a direct comparison of the methodological advantages and shortcomings of MR.

Together, this work establishes microrespirometry-based inhibitor screening assays as a rapid, sensitive, and versatile platform for disentangling the efficacy, specificity, and mode of action of nitrification inhibitors. By providing high-resolution, real-time insights into microbial respiration, this approach advances our ability to mechanistically evaluate novel candidate nitrification inhibitors for sustainable N management. Ultimately, such tools are essential to support the development of targeted strategies to mitigate agricultural N losses.

## Materials and Methods

### Nitrification inhibitors

The synthetic nitrification inhibitors NP (Sigma Aldrich; >98%), DCD (Sigma Aldrich; 99%), and PA (Sigma Aldrich; 98%), and the organic solvent dimethyl sulfoxide (DMSO; Sigma Aldrich; >99.9% / Carl Roth; >99.8%) were used in this study. Inhibitor solutions were freshly prepared in DMSO or sterilized water (DCD in MR experiments) before use.

### Organisms and cultivation

Pure cultures of *Nitrosomonas europaea* Nm50 (AOB), *Nitrosospira multiformis* NI2 (AOB), *Nitrospira inopinata* (comammox), and *Nitrospira moscoviensis* (NOB) were grown as batch liquid cultures in mineral media, in the dark, at their respective optimal growth temperature (28 or 37°C) and pH (7.8) (Table S1). AOB and comammox culture medium contained (L^-1^): 50 mg KH_2_PO_4_, 75 mg KCl, 24 mg MgSO_4_ x 7 H_2_O, 584 mg NaCl, 1 g HEPES, 1 mL TES (trace element solution) and 1 mL SWS (selenium-wolfram solution) (Jung et al., 2022). *N. moscoviensis* was grown in medium containing (L^-1^): 10 mg CaCO_3_, 500 mg NaCl, 50 mg of MgSO_4_ × 7H_2_O, 150 mg KH_2_PO_4_, 10 mg NH_4_Cl, 1 mL TES and 1 mL SWS. In all cases, the pH of the growth medium was adjusted to 7.8 with NaOH prior to autoclaving. Substrate (NH_4_Cl or NaNO_2_, 1-4 mmol L^-1^) was supplemented from sterile stock solutions as needed and the pH was maintained at 7.8 by addition of sterile NaHCO_3_.

### Microrespirometry assays

For MR experiments, biomass was harvested from substrate replete cultures and concentrated by centrifugation (3,000-4,500 g, 20°C, 15-20 min) in 10 kDa-cutoff, Amicon Ultra-15 centrifugal filter units (Merck Millipore, Germany). Concentrated biomass was washed with and resuspended in sterile substrate-free media, before being incubated for at least 30 min in a recirculating water bath set to the optimum growth temperature of the culture. Once acclimatized, the biomass was transferred to MR chambers.

Whole-cell respiration rates were determined in an MR system equipped with a 4-channel microOptode meter (Opto-F4 UniAmp) and O_2_ MicroOptodes (50 µm tip diameter, Unisense Denmark). Glass MR chambers (∼2 mL) containing glass-coated magnetic stirring bars were filled headspace-free with concentrated culture biomass, sealed with MR injection lids, and submerged on a magnetic stirrer rack (300 rpm) in a recirculating water bath set to the optimal growth temperature for each culture. An O_2_ MicroOptode was inserted into each MR chamber and the background cellular respiration was recorded for at least 30 minutes. After a stable background respiration rate was achieved, up to three substrate injections (NH_4_Cl or NaNO_2_) were performed with a Hamilton gas-tight syringe to achieve a maximum respiration rate in the chamber. Initial substrate concentrations were chosen for each culture-based on previously described whole-cell kinetic measurements (Jiang and Bakken, 1999; Nowka et al., 2015; Kits et al., 2017; Jung et al., 2022; Table S1). After the initial substrate injections and each subsequent injection of DMSO, inhibitor, or substrate, the oxygen consumption rate was recorded for 3-5 min. In individual MR chambers, substrate, DMSO and/or inhibitors were injected multiple times, increasing their concentration in the MR chamber over time.

NH_4_Cl or NaNO_2_ were provided as the initial substrate when determining inhibitor efficacy and organism specificity. If complete inhibition was observed during the experiment, multiple substrate (NH_4_Cl) injections were performed after the last inhibitor injection to assess inhibitor competitiveness. Thereafter, in a subset of experiments performed with *N. inopinata* N_2_H_4_ was injected to test for the enzyme specificity of each inhibitor.

In experiments performed specifically to assess up to which concentration inhibitors act in an AMO specific manner, N_2_H_4_ (300 µmol L^-1^, Table S1) was provided as the initial substrate, followed by injections of inhibitors until an inhibitor concentration many fold exceeding the respective inhibitors IC_100_ for AMO was reached. In order to account for any potential DMSO-driven inhibition due to the large quantities of DMSO injected in these MR assays, additional DMSO controls for each biomass preparation were performed. Here, harvested, concentrated, and acclimatized biomass from a single culture was split into two MR chambers. Injections of substrate, and then inhibitor dissolved in DMSO or DMSO only, were performed in parallel. The relative activity (%) at each inhibitor concentration was then corrected for any observed DMSO induced inhibition. A schematic description of the experimental workflow and possible MR activity observations is shown in Figure 1.

**Figure 1.**
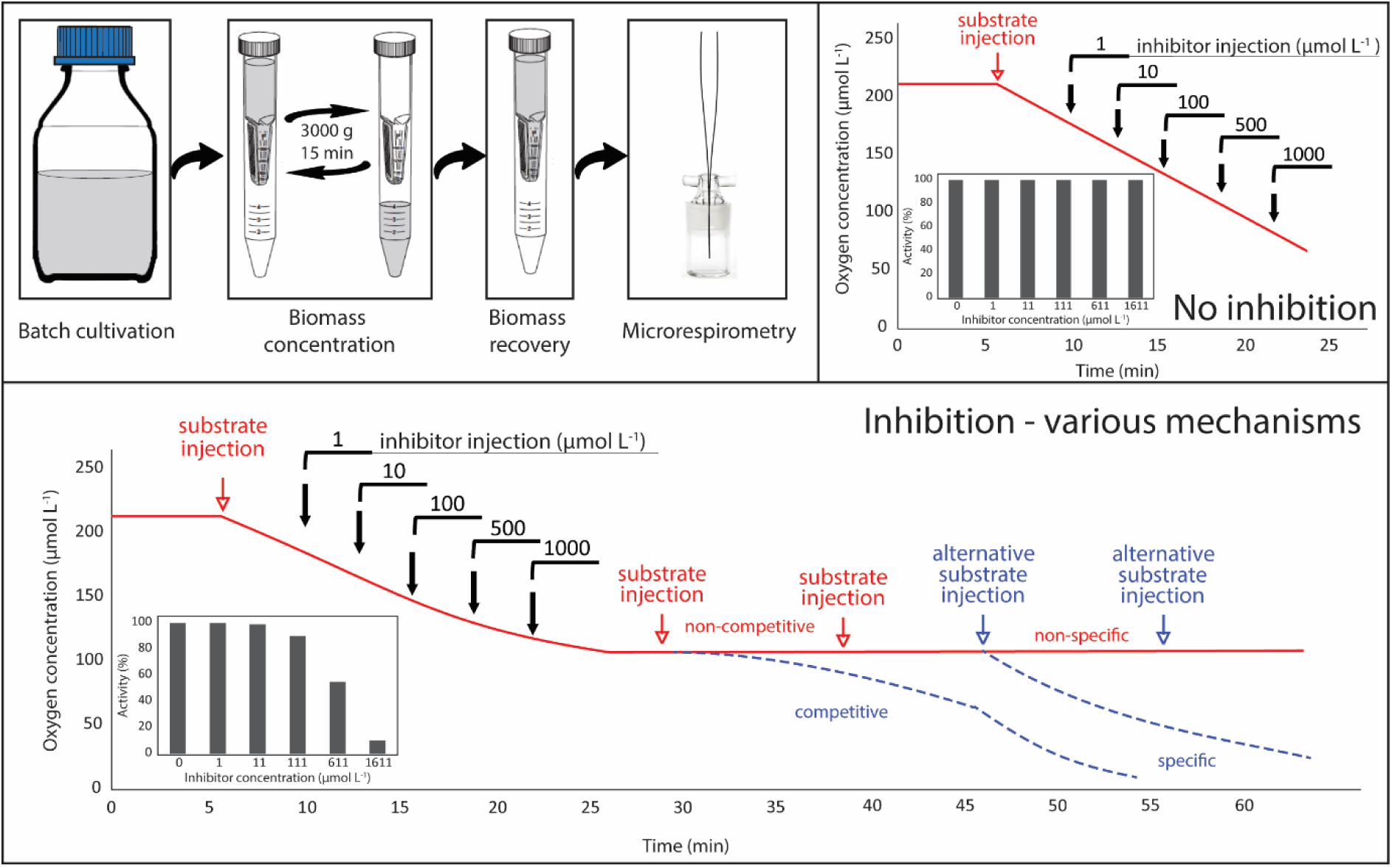
Microrespiration-based workflow schematic. Upper panel left side: Culture preparation workflow for candidate inhibitor microrespirometry experiments. **Upper panel right side:** Example oxygen consumption curves and percent activity profiles obtained by microrespirometry of a non-inhibitory compound. **Lower panel:** Example oxygen consumption curves and percent activity profiles obtained by microrespirometry of an inhibitory compound. The response of a non-competitive (solid red line) versus competitive (dashed blue line) inhibitory compound to an additional substrate injection, as well as the response of a enzyme specific (dashed blue lines) and non-specific (solid red line) inhibitor to injection of an alternative substrate is also shown.

### Growth assays

In growth assays, inhibitor efficacy was determined by providing each inhibitor at 4-6 different concentrations (Table S2) to four replicate cultures (50 mL) per inhibitor concentration for each ammonia-oxidizing microorganism. A dilution series of inhibitor stocks was prepared in pure DMSO so that the final concentrations of DMSO were 0.1% v/v in all incubations. Control incubations, supplemented with 0.1% v/v DMSO only, were run in parallel. All cultures were subsampled every 2-3 days until the control incubations were substrate depleted (1 mmol L^-1^). Subsamples were stored at -20°C and used to determine NO_2_ ^-^ and NO_3_ ^-^ concentrations with an acid VCl_3_/Griess assay (Miranda et al., 2001).

### Data Analysis and Statistics

Oxygen consumption rates (i.e., respiration rates, corrected for background respiration) in the presence of increasing concentrations of DMSO, PA, NP, and DCD were determined and used to calculate the inhibitor/solvent concentration-dependent reduction in respiration rates. These are expressed as percent residual activity at a given inhibitor concentration. Inhibitor efficacy was assessed by determining the half maximal inhibitory concentration (IC_50_). The IC_50_ is defined as the concentration of an inhibitor that reduces the respiration rate coupled to ammonia oxidation activity of each species tested to 50% in an MR assay, or as the concentration of an inhibitor that reduces the nitrite/nitrate production of each species tested to 50% in a batch growth assay.

For MR assays, the oxygen consumption rate in the absence of inhibitors was taken as 100% activity. IC_50_ values for each species-inhibitor pair were modeled from dose-response curves of relative respiration rates and inhibitor concentrations (corrected for the inhibitory effect of DMSO in *N. europaea* MR assays - described below) using a logistic regression model (i.e., the Hill-Langmuir equation, Sebaugh, 2010, Equation 1):

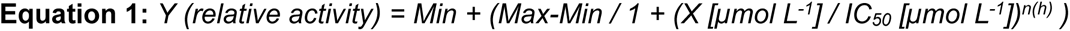

where Y is the observed relative respiration rate at inhibitor concentration X, Max is the maximum relative activity, Min is the minimum relative activity, IC_50_ is the inflection point of the modeled sigmoid dose-response curve, and n(h) is the Hill coefficient. Since relative activity data was used to determine the IC_50_, Min was set to 0 and Max to 1, restraining the logistic regression model to two parameters.

When NH ^+^ was provided as a substrate, a significant inhibitory effect of DMSO on the respiratory activity of *N. europaea* was determined by a paired t-test and was found to be linearly dependent on DMSO concentration (Figure 2). To account for this DMSO inhibition the respiration rate upon inhibitor addition was corrected for the expected inhibition at the corresponding DMSO concentration using the following equations (Equation 2-4):

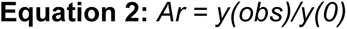

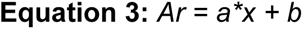

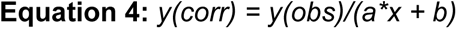

where y(0) is the uninhibited oxidation rate, y(obs) is the observed oxidation rate after DMSO injection, y(corr) is the corrected oxidation rate in µmol O_2_ h^-1^, x is the DMSO concentration in µmol L^-1^, Ar is the relative activity/residual activity, *a* is the slope, and *b* is the intercept of Ar expressed as a linear function of x.

**Figure 2.**
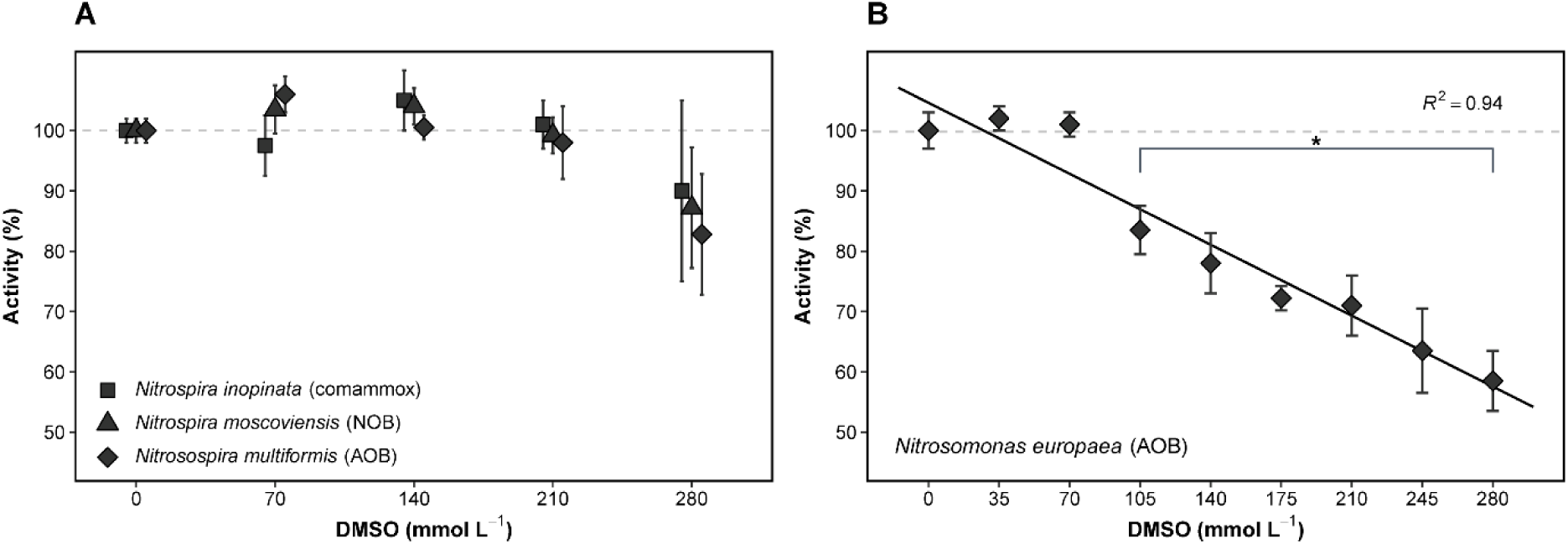
Inhibitory effect of DMSO on nitrifier cellular respiration. Percent cellular respiration activity with multiple additive DMSO injections (35 mmol L^-1^, 0.25% v/v) up to a final concentration of 280 mmol L^-1^ or 2% v/v. The error bars represent the standard deviation of three independent culture replicates. The asterisk indicates the onset of significant inhibition as determined by a paired t-test (p< 0.05). **(A)** No significant effect of DMSO on cellular respiration was observed for *N. multiformis*, *N. inopinata* or *N. moscoviensis*. **(B)** A linear trendline was used to model DMSO concentration dependent inhibition in *N. europaea*.

When inhibitor specificity was tested, a paired t-test was used to determine the significance of non-enzyme specific inhibition. Here, the respiration rate of a non-inhibited N_2_H_4_-oxidizing culture was compared to its respiration rate at a given inhibitor concentration, after correcting for the effect of DMSO alone.

For growth assays, the average reaction product concentration (NO_2_^-^ or NO_2_^-^ + NO_3_^-^; for the AOB or comammox organisms, respectively) observed in DMSO-amended control incubations at a given time point was taken as 100% activity. Relative activity values at given initial inhibitor concentrations across all sample timepoints were then calculated. IC_50_ values for each species-inhibitor pair were modeled from dose-response curves of relative product concentrations and inhibitor concentrations using the same logistic regression model as described above (Equation 1), adapted to a product concentration-based evaluation so that Y was the observed relative product concentration at inhibitor concentration X.

To statistically assess if individual inhibitors exhibited uniform potency shifts between assays, Pearson correlation coefficients (*r*) were calculated between the log_10_- transformed IC_50_ values obtained from MR-based and growth-based assays across all three ammonia-oxidizing species. A paired t-test was used to evaluate the statistical significance of the log_10_-transformed Pearson correlation coefficients.

## Results and Discussion

### Effect of DMSO on respiration activity in MR experiments

DMSO is a widely used cryopreservant and organic solvent. PA and NP, like many other nitrification inhibitors, are poorly water soluble and routinely dissolved in DMSO for laboratory testing. Therefore, the effect of DMSO on nitrifier activity with their primary substrate (ammonium or nitrite) in the MR setup was tested. No significant effect of DMSO on the maximum respiration rate of *N. inopinata*, *N. multiformis*, or *N. moscoviensis* was observed up to a concentration of ∼280 mmol L^-1^ DMSO (∼2% v/v) (Figure 2). In contrast, the respiration rate of *N. europaea* showed higher sensitivity and linearly decreased with DMSO concentration, starting at ∼105 mmol L^-1^ (0.75% v/v). At ∼2% v/v DMSO, the respiration rate of *N. europaea* was reduced by 43 ±5% (Figure 2).

To account for the inhibitory effect of DMSO on *N. europaea*, the total inhibitory effect of inhibitors dissolved in DMSO (PA and NP) was corrected for the inhibitory effect of the total DMSO chamber concentration (see *Methods*), assuming an additive inhibitory effect of DMSO and the respective inhibitor. Likewise, in inhibitor specificity assays with N_2_H_4_ as substrate, where DMSO concentrations at the end of MR experiments were expected to exceed 2% due to the high number of inhibitor injections performed, DMSO control MR-assays were performed in parallel with biomass from the same harvest. Where necessary, the effect of inhibitors was corrected for the inhibitory effect of DMSO alone. Finally, to eliminate the possibility of DMSO interference with efficacy results in growth assays, inhibitor efficacy was directly compared against a DMSO only control.

In literature, the effect of DMSO on nitrifiers and nitrification activity is debated. In several high cell density substrate conversion assay studies, no effect of DMSO on nitrite production rates by various AOA and AOB, including *N. europaea* and *N. multiformis*, was observed (Wright et al., 2020 Zhao et al., 2020a; Papadopoulou et al., 2020; Kaur-Bhambra et al., 2022). However, other studies have reported reduced ammonia oxidation rates in the presence of 0.1% v/v DMSO for the AOA isolates *N. viennensis* (Kaur-Bhambra et al., 2022) and *N. maritimus* (Sauder et al., 2016), as well as for *N. europaea* (Wright, 2021). Likewise, a negative effect of DMSO additions on nitrification rates in environmental samples has been reported (Saari and Martikainen, 2001; Lehtovirta-Morley et al., 2013). Notably, the DMSO concentration at which we observed inhibition was higher than analyzed in most other studies (Wright, 2021., Zhao et al., 2020a; Papadopoulou et al., 2020). Overall, based on observations from the current study as well as a literature review, different species, different strains of the same species, and likely even the same ammonia-oxidizing strains under different experimental conditions, display variable levels of sensitivity to DMSO. Our results thus underline the necessity of carefully assessing and accounting for the effect of any organic solvents in nitrification inhibitor efficiency studies.

### Phenylacetylene (PA) mediated inhibition in microrespiration and growth assays

PA is a well characterized, specific, non-competitive inhibitor of AMO in AOB (Hyman and Wood, 1985; Vannelli et al., 1996). More recently PA has also been shown to have a specific and irreversible inhibitory effect on ammonia oxidation by the AOA isolate *Ca.* Nitrosocosmicus franklandianus (formerly *Ca.* Nitrosocosmicus franklandus), although higher concentrations (>10 µmol L^-1^) are required to achieve inhibition (Wright et al., 2020). While PA is less commonly used in environmental studies (e.g. Im et al., 2011; Bayer et al., 2025), and not routinely applied in agriculture, PA was selected as a proof-of-concept inhibitory compound, due to its well characterized mode of inhibition (Hyman and Wood, 1985; Lontoh et al., 2008; Wright et al., 2020).

In MR assays, complete inhibition of ammonia oxidation was achieved at PA concentrations of ∼10 µmol L^-1^, while the IC_50_ was <1 µmol L^-1^ PA for all three ammonia-oxidizing species tested. *N. europaea* was the most sensitive to PA, with an IC_50_ of ∼70 nmol L^-1^, while the IC_50_ values of *N. inopinata* and *N. multiformis* were 2-13-fold higher (0.14 and 0.94 µmol L^-1^ respectively; Figure 3).

**Figure 3.**
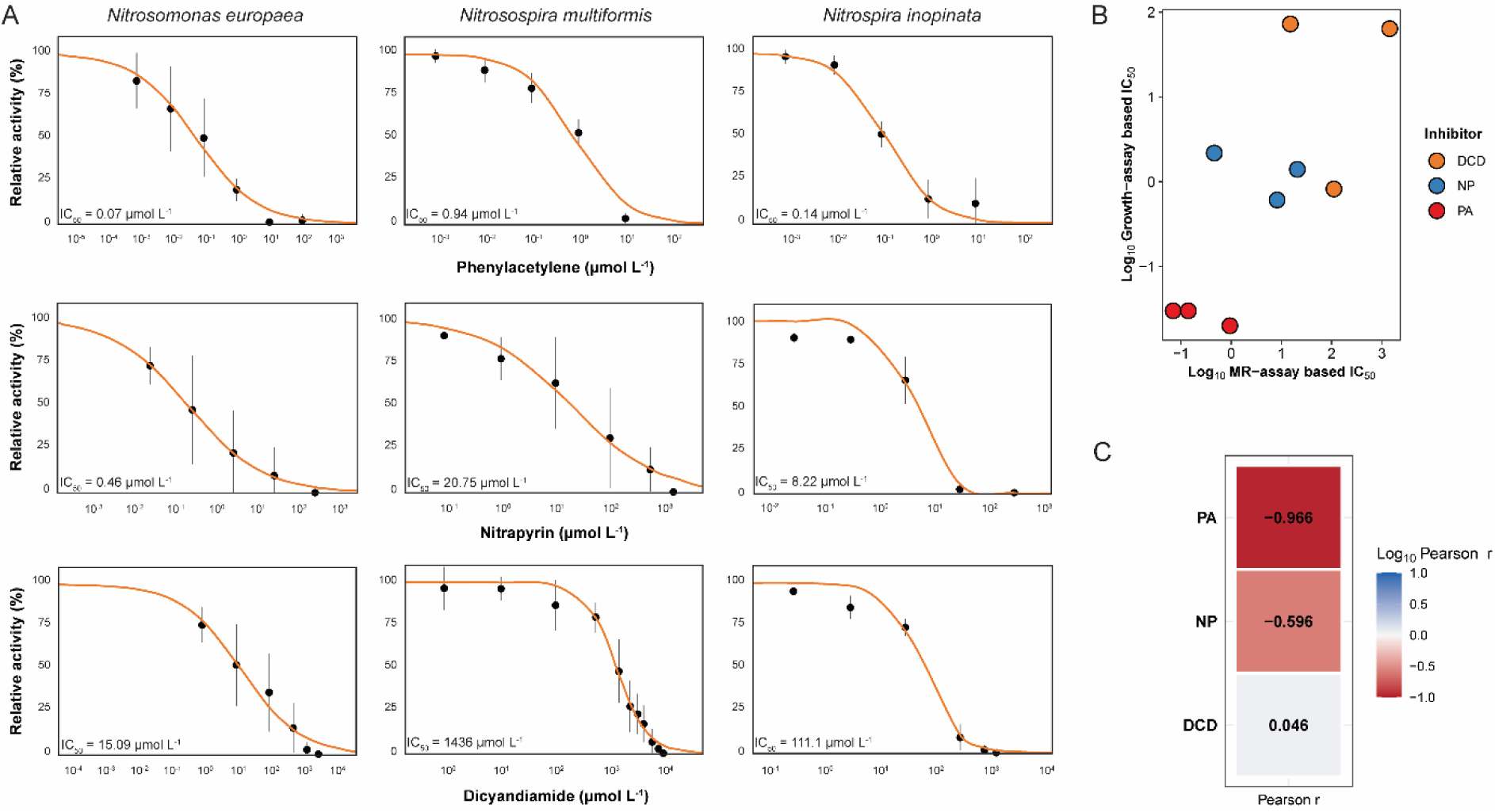
**A** Constrained logistic regression models of respiration inhibition when ammonium was provided as the sole substrate. Relative respiratory activity of *N. europaea* (left), *N. multiformis* (middle) and *N. inopinata* (right) cultures in response to increasing concentrations of phenylacetylene (top), nitrapyrin (middle) and dicyandiamide (bottom). IC_50_ values were modeled from dose-response curves using a logistic regression model (i.e., the Hill-Langmuir equation, Sebaugh, 2010). Error bars represent the standard deviation of three independent culture replicates. **B** Scatter plot of log_10_-transformed IC_50_ values derived from MR-based inhibitor efficacy assays (MR, x-axis) versus growth-based inhibitor efficacy assays (G, y-axis). **C** Paired Pearson correlation coefficients (r) between log_10_-transformed IC_50_ values derived from MR-based inhibitor efficacy assays versus growth-based inhibitor efficacy assays for each of the three inhibitors. None of the reported correlations reached statistical significance.

In growth assays complete inhibition of ammonia oxidation was achieved at PA concentrations of ∼1 µmol L^-1^ for all three tested species, which is in line with previous reports from similar experiments (Hyman and Wood, 1985; Vannelli et al., 1996; Lontoh et al., 2008). In all cases, the IC_50_ values determined in growth assays were lower than those determined in MR assays (Figure 3, Figure S1). For *N. europaea* and *N. inopinata*, the growth assay-based IC_50_ of PA was comparable (only ∼2.5 to 4-fold lower, at ∼20 nmol L^-1^ and ∼30 nmol L^-1^; respectively) than determined by MR. Interestingly, *N. multiformis* was substantially more sensitive to inhibition by PA in the growth assay, with an IC_50_ value nearly 40-fold lower (∼30 nmol L^-1^) than the IC_50_ of ∼940 nmol L^-1^ observed in MR assays (Figure 3, Figure S1). The MR- and growth-assay IC_50_ values were strongly, but not significantly (p=0.17) correlated (Figure 3), likely due to the small sample size.

### Nitrapyrin (NP) mediated inhibition in microrespiration and growth assays

NP has been investigated as a nitrification inhibitor since the late 1950s (Lees, 1954; Goring et al., 1962a; 1962b), and is the most widely commercially used agricultural nitrification inhibitor (Subbarao et al., 2006a, Coskun et al., 2017). Numerous studies have demonstrated that NP inhibits ammonia oxidation in both *Nitrosomonas*- and *Nitrosospira*-related AOB (e.g., Campbell and Aleem, 1965; Hooper and Terry, 1973; Belser and Schmidt, 1981; Shen et al., 2013; Papadopoulou et al., 2020), as well as in all AOA strains tested to date (Lehtovirta-Morley et al., 2013; Shen et al., 2013; Papadopoulou et al., 2020). However, prior to this study, NP inhibition in a pure comammox *Nitrospira* culture had not been tested. Notably, the reported inhibitory concentrations for NP vary widely- from less than 1 to over 200 µmol L^-1^, for the same AOB species (*N. multiformis*) across studies (e.g. Shen et al., 2013; Papadopoulou et al., 2020). Furthermore, there have been conflicting results on the inhibitory effect of NP on ammonia oxidation by comammox *Nitrospira* (e.g. Li et al., 2020; Papadopoulou et al., 2022; Zhou et al., 2020b).

In MR-assays, ammonia oxidation by *N. inopinata* and *N. europaea* was completely inhibited by ∼50 µmol L^-1^ NP, whereas *N. multiformis* was less sensitive, requiring concentrations of >500 µmol L^-1^ to achieve complete inhibition (Figure 3). This was also reflected in the IC_50_ values with *N. europaea* (0.46 µmol L^-1^) and *N. inopinata* (8.22 µmol L^-1^) being more sensitive than *N. multiformis* (20.75 µmol L^-1^) for NP (Figure 3).

In contrast to the different levels of sensitivity observed in the MR-assays, in growth assays, all three ammonia-oxidizing species responded very similarly. Total inhibition was always achieved at <10 µmol L^-1^ NP (Figure S1), equaling 5- to >50-fold lower concentrations than needed in MR-assays. Interestingly, IC_50_ values for NP in growth assays were an order of magnitude lower than in MR-assays for *N. inopinata* (0.61 µmol L^-1^) and *N. multiformis* (1.4 µmol L^-1^), but about five times higher for *N. europaea* (2.18 µmol L^-1^) (Figure 3, Figure S1). No clear correlation between MR- and growth-assay based IC_50_ values for NP was observed (Figure 3).

### Dicyanamide (DCD) mediated inhibition in microrespiration and growth assays

DCD, like NP, is a well-known and widely used nitrification inhibitor in agriculture (Prasad et al., 1971; Slangen and Kerkhoff, 1984; Coskun et al., 2017). In general, required DCD concentrations are reported to be much higher (>500 µmol L^-1^) than for NP or PA (Shen et al., 2013; O’Sullivan et al., 2017; Papadopoulou et al., 2020).

As already observed for NP, *N. inopinata* and *N. europaea* were more sensitive to inhibition by DCD in MR assays than *N. multiformis*. Total inhibition of respiration in *N. inopinata* and *N. europaea* was observed at concentrations of ∼1.5 mmol L^-1^ DCD, while complete inhibition of *N. multiformis* was only observed at concentrations exceeding 10 mmol L^-1^ DCD (Figure 3). A comparatively low IC_50_ value for DCD was determined for *N. europaea* (15.09 µmol L^-1^), whereas an extremely high IC_50_ value was observed for *N. multiformis* (∼1.5 mmol L^-1^). The IC_50_ DCD value of *N. inopinata* in MR-assays was, like already observed for PA and NP, between those observed for the two AOB strains, at 111.1 µmol L^-1^ (Figure 3).

In growth assays with DCD, total inhibition was again, as with PA and NP reached with lower concentrations than in MR-assays: both *N. europaea* and *N. multiformis* were fully inhibited with 500 µmol L^-1^ DCD and total inhibition for *N. inopinata* was achieved at 100 µmol L^-1^ (Figure S1). Interestingly, as observed with NP, the growth assay-based IC_50_ for DCD in *N. inopinata* and *N. multiformis* was much lower than the MR assay-based IC_50_ (>100x and >20x, respectively) (Table 1); but *N. europaea*, again had a higher IC_50_ for DCD in growth assays (72.7 µmol L^-1^, Figure S1) than in MR assays (15.09 µmol L^-1^; Figure 3). Again, no clear correlation between MR- and growth-assay based IC_50_ values for DCD was observed (Figure 3).

### Determining enzyme specificity of inhibitors through MR assays

Assays or workflows which allow for a simple and reliable investigation of inhibitor specificity are essential for inhibitor characterization. MR assay-based experiments provide an intrinsically simple way to determine the enzyme specificity of potential inhibitors, since the same setup and readout can be used to assess the effect of inhibitors on AMO-dependent respiration, AMO-independent respiration, and the respiration of non-ammonia-oxidizing microorganisms.

Previous research on the specificity of PA, NP, and DCD report all three inhibitors are non-competitive and at least partially AMO specific (Campbell and Aleem, 1965; Hyman and Wood, 1985; Zacherl and Amberger, 1990; Vannelli et al., 1996; Wright et al., 2020). While PA is a mechanics-based suicide inhibitor (Wright et al., 2020), the inhibitory mechanisms and enzyme specificity of NP and DCD are not as clear. The first physiological study investigating the inhibition mechanism of NP (Campbell and Aleem, 1965) and DCD (Zacherl and Amberger, 1990) in *N. europaea* described AMO specific inhibition, as hydroxylamine oxidation remained essentially unaffected by NP and DCD concentrations which led to a full inhibition of ammonia oxidation activity. Further evidence for a mechanistic inhibition of ammonia oxidation by NP is derived from the fact that it can be co-metabolized by AMO in *N. europaea* (Hooper and Terry, 1973; Vannelli and Hooper, 1992; Vannelli and Hooper, 1993). Yet, the primarily inhibitory mechanism of NP is nevertheless considered to be based on the chelation of copper, a metal cofactor essential for AMO functioning (Campbell and Aleem, 1965). DCD, likewise, presumably also primarily acts via copper chelation (McCarty, 1999). Copper is not only the active site metal of AMO, but also an important cofactor for a number of other microbial enzymes (e.g. cytochrome *c* oxidase, nitric and nitrous oxide reductase, some superoxide dismutases; Argüello et al., 2013). Thus, a copper-chelating inhibitor can induce a variety of non-enzyme-specific effects in ammonia-oxidizers and other off-target microorganisms.

We used a MR assay to test inhibitor specificity in two ways:

1. MR-assays with a non-target microorganism were performed to test if the three inhibitors act specifically on ammonia-oxidizing microorganisms, or if inhibition was conferred by other mechanisms such as cytotoxicity and copper chelation. For this, the NOB *N. moscoviensis*, which is phylogenetically closely related to the comammox strain *N. inopinata,* was used. The rate of nitrite oxidation by *N. moscoviensis* was not significantly affected by DCD and PA at concentrations needed to fully inhibit ammonia oxidation (Figure S2), confirming that the inhibition observed was specific to ammonia-oxidizing microorganisms (Figure 3). At a NP concentration of ∼IC_100_ for the three ammonia-oxidizing cultures respiration of *N. moscoviensis* showed slight inhibition, indicating the mechanism(s) of inhibition is not specific to ammonia-oxidizing microorganisms at elevated concentrations (Figure S2).
2. The effect of inhibitors on AMO-independent respiration by ammonia-oxidizing microorganisms was tested by supplying N_2_H_4_ as an AMO-activity bypassing substrate in the MR-assay setup (Figure 4). In agreement with literature (Hyman and Wood, 1985; Wright et al., 2020), no statistically significant inhibition (p>0.05) of AMO-independent N_2_H_4_-driven respiration was observed in the three analyzed ammonia-oxidizing species, even in the presence of PA concentrations three orders of magnitude higher than those needed to fully inhibit ammonia oxidation (Figure 4). Notably, these results also support the hypothesis that PA, like other alkynes (Sakoula et al., 2022), is covalently bound by the structurally yet uncharacterized AMO of comammox *Nitrospira*, thus acting as a suicide inhibitor as previously described for the related AOB AMO enzymes (Hyman and Wood, 1985; Vannelli et al., 1996). However, NP and DCD mediated inhibition was not consistently AMO specific. At high NP concentrations, exceeding 5.5 µmol L^-1^ for *N. inopinata* and 550 µmol L^-1^ for *N. multiformis*, a significant (p<0.01) inhibitory effect was observed on N_2_H_4_-driven respiration. Notably, this was not the case for *N. europaea* (Figure 4, p>0.05), where in agreement with previous literature reports NP did act as a AMO specific inhibitor at all tested concentrations (Campbell and Aleem, 1965). The fact that NP induced inhibition is only partially enzyme specific may help explain the inconsistencies reported in literature across cultures and environmental samples for the efficacy and mode of action of NP. Finally, increasing DCD concentrations led to increasing levels of inhibition of AMO-independent N_2_H_4_-driven respiration in all three ammonia-oxidizing cultures (Figure 4), but highly significant inhibition (p<0.01) was only observed at concentrations around or above the IC_100_ for the respective species. As the primary mechanism of action for DCD is assumed to be based on copper chelation, our results indicate that the role of copper, beyond being the AMO active site metal in ammonia-oxidizing microorganisms, might be linked to AMO-independent susceptibility to DCD.

**Figure 4.**
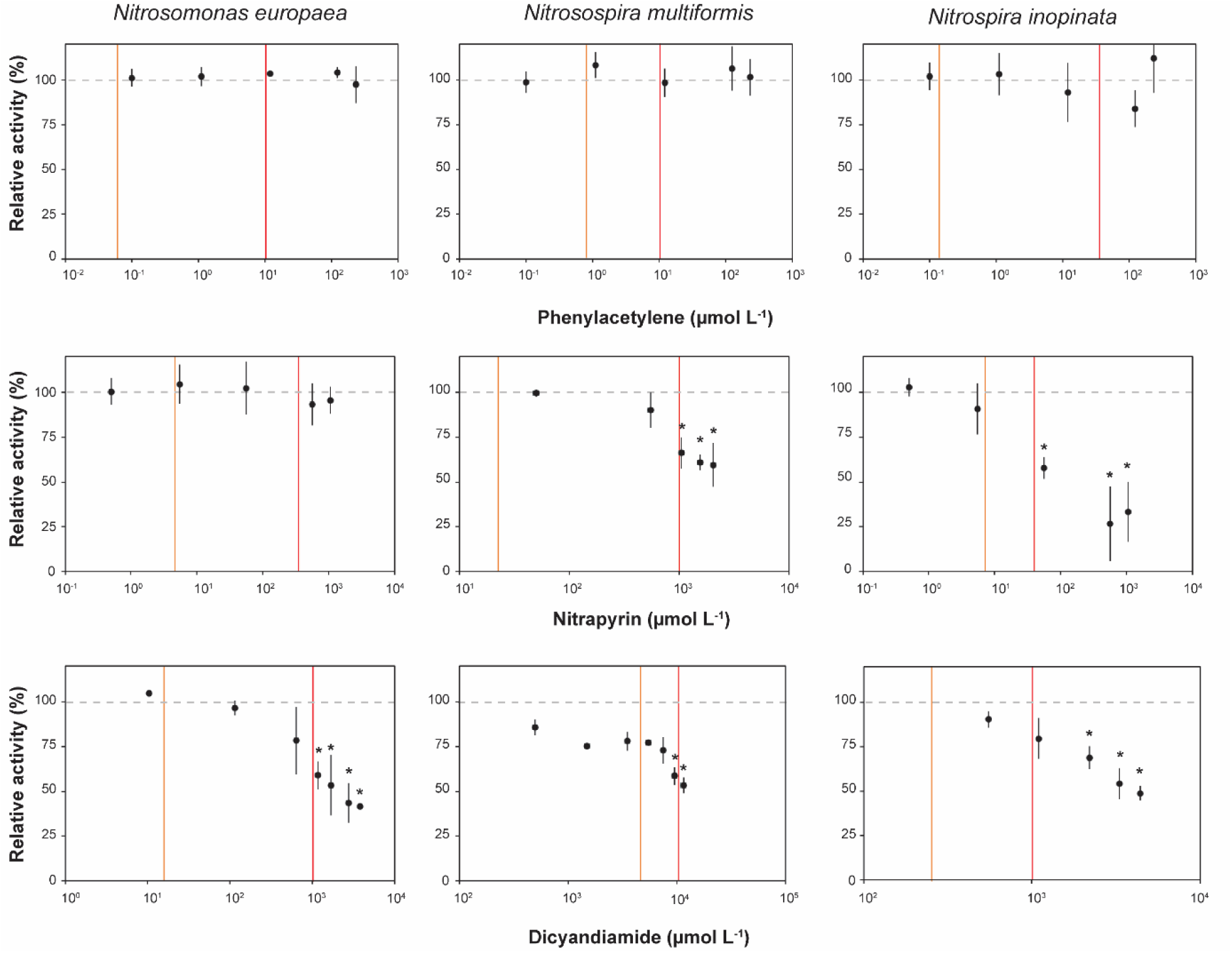
MR-assay based determination of inhibitor AMO-specificity. Relative respiratory activity of *N. europaea* (left), *N. multiformis* (middle) and *N. inopinata* (right) cultures in response to multiple successive injections of phenylacetylene (top), nitrapyrin (middle) and dicyandiamide (bottom) during AMO-independent N_2_H_4_-driven respiration. Error bars represent the standard deviation of three independent culture replicates. Asterisks indicate significant inhibition as determined by a paired t-test (p< 0.01). The IC_50_ values for MR assay-based AMO inhibition for the same culture-inhibitor combination are marked with the orange line, and for IC_100_ values with the red line.

### Determining inhibitor efficacy, competitiveness and specificity in a single MR-assay

After establishing and evaluating the different aspects (solvent effect, efficacy, and specificity assays) of MR-assay suitability for mechanistic inhibitor characterization, we combined them into an MR-assay in which inhibitor efficacy, competitiveness, and enzyme specificity are assessed in a single MR-chamber, using the same biomass sample (Figure 5). This assay approach was performed with all three inhibitors and the comammox isolate *N. inopinata*, as inhibition of this species by PA, NP and DCD had not been profiled before this study and further mechanistic insights were warranted. Here, NH_4_Cl was provided as primary substrate to determine baseline 100% respiration activity, followed by multiple inhibitor injections to determine inhibitor efficacy (data included in Figure 3). Once total inhibition was achieved, the competitiveness of the inhibitor was tested with multiple additional injections of NH_4_Cl (as detailed in Figure 1). PA, NP, and DCD were all determined to inhibit *N. inopinata* in a non-competitive manner (Figure 5). Subsequently, the AMO-bypassing substrate N_2_H_4_ was injected into the same MR chamber. N_2_H_4_ injections in all cases restored *N. inopinata* respiration activity (Figure 5), with the fraction of activity restored being high for PA and DCD and low for NP, in accordance with the previously determined degree of off-target inhibition at ∼IC_100_ for the respective inhibitors (Figure 4).

**Figure 5.**
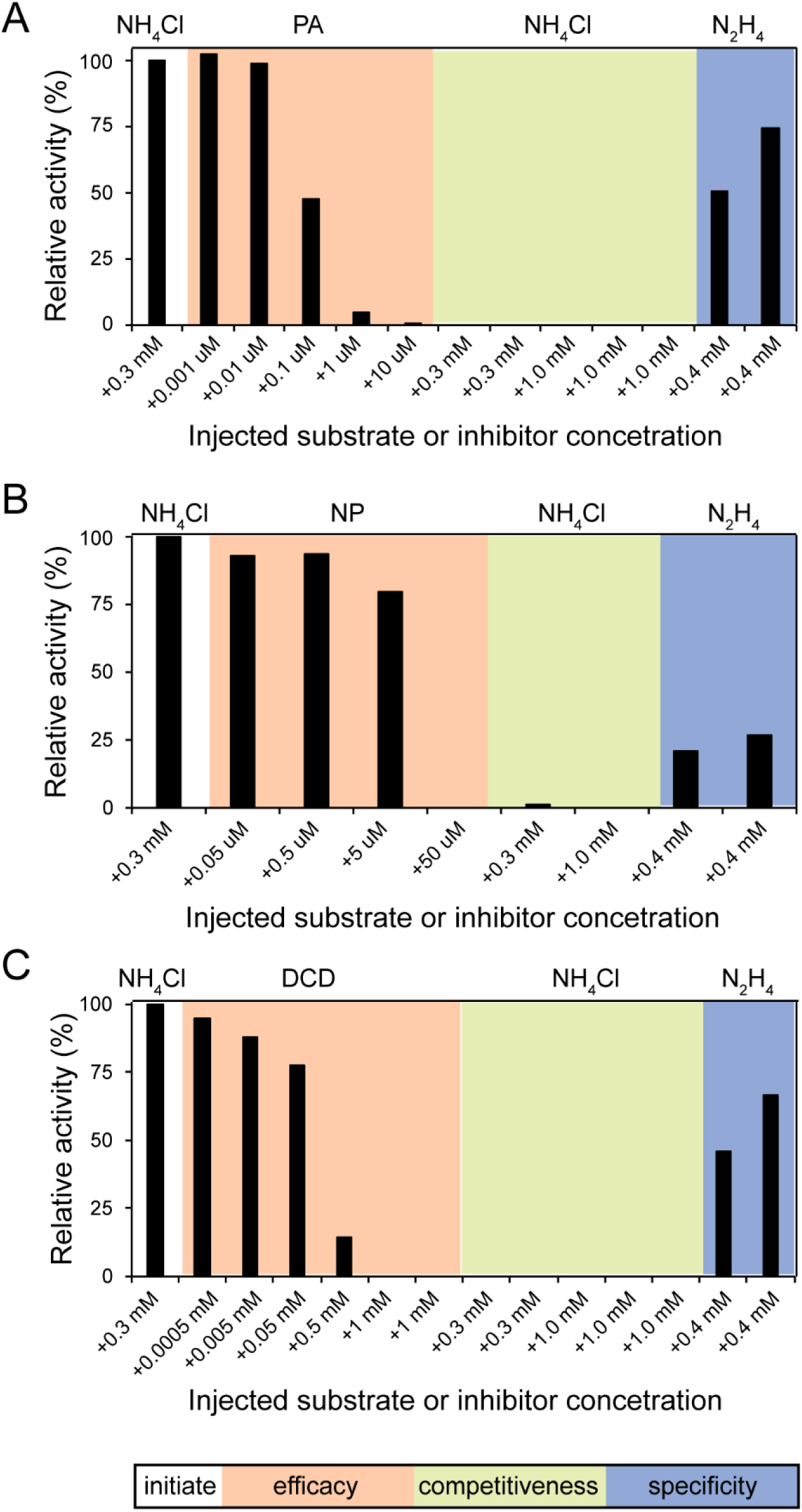
A single chamber MR-based approach to determine inhibitor efficacy, competitiveness, and enzyme specificity. Relative activity of *N. inopinata* respiration in the presence of different concentrations of phenylacetylene (**A**), nitrapyrin (**B**) and dicyandiamide (**C**). The initial injection in all cases was NH_4_Cl (0.3 mM), followed by multiple injections of the inhibitor (efficacy testing), NH_4_Cl (competitiveness testing), and N_2_H_4_ (specificity testing), in that order. X-axis values represent the concentration injected into the chamber (individual injection amounts) and not the total concentration of the inhibitor, NH_4_Cl, or N_2_H_4_ in the chamber (additive amounts). N_2_H_4_ relative activity rates were multiplied by 1.5x, adjusting for expected maximal oxygen consumption rates. All data from each panel represents a single MR chamber experiment and is therefore an independent culture replicate.

## Conclusions

Using the well-studied proof-of-concept nitrification inhibitors PA, NP and DCD, we demonstrate how MR-assays provide a rapid, convenient, and versatile methodological toolkit for the mechanistic investigation of potential novel nitrification inhibitors (Figure 1). With this MR-workflow, we successfully reproduced previously published patterns of inhibitory specificity and mode of action in AOB. Moreover, we provided the first pure culture insights into how PA, NP, and DCD inhibit ammonia oxidation by the comammox isolate *N. inopinata* (Figure 5).

Notably, we observed that inhibitor IC_50_ values are assay-method dependent, with results for DCD differing up to two orders of magnitude between MR- and growth-based assays, and no correlation between results observed for DCD and NP (Figure 3). Notably, IC_50_ values for PA were strongly, but not significantly correlated between the methods (Figure 3). This high degree of discrepancy for NP and DCD, when experiments are performed in the same laboratory with the same strains may help explain the substantial variability in reported efficacy values for nitrification inhibitors across studies. These differences are likely in part due to the not fully enzyme specific inhibition mechanism of DCD and NP (Figure S2, Figure 4, Figure 5), and also likely influenced by the fundamentally different physiological processes captured by the two different methods. Growth, substrate conversion, and respiration integrate different aspects of cellular activity under inhibitor stress and therefore cannot be expected to yield identical quantitative outcomes for inhibitors that exert some degree of off-target (i.e. non-enzyme-specific) effect. Other methodological factors likely also contribute to variability. For instance, cell density is rarely standardized or reported across studies. In our experiments, biomass from late exponential-phase was concentrated 10- to 100-fold for MR assays (Table S2), whereas for growth assays biomass washed and diluted approximately 5-fold, resulting in differences in biomass density of up to two orders of magnitude between approaches. However, inhibitor efficiency does not scale proportionally with these differences, highlighting the non-linear relationship between biomass concentration and apparent inhibitor efficiency. Experimental duration (minutes in MR-assays, minutes to hours in substrate conversion assays, and days to weeks in growth assays), as well as culture physiological state and fitness, likely also strongly influence the resulting inhibitor response profiles.

Taken together, these observations suggest that characterizing the specificity and mode of action of an inhibitor might be more critical and informative than aiming to define a single, universal inhibitory dose. Instead, inhibitor efficacy should be interpreted in context-dependent orders-of-magnitude range. This perspective is particularly important given that in natural environments, like soils, inhibitor concentration is unlikely to be homogeneous in either space or time. Within this framework, MR-based assays provide a particularly valuable approach, as they enable a readout of efficacy, specificity, competitiveness, and reversibility (albeit not tested here), using the same physiological trait (i.e. respiration) from a single biomass sample. In addition, they are the fastest of the available approaches, and provide a continuous real-time readout, which enables on the fly experimental flexibility. This integrated capability makes MR a powerful addition to the growing methodological toolbox for studying compound–microbe interactions, for nitrification inhibitors and beyond.

## Acknowledgement

We would like to thank Johanna Palmetzhofer for support with culture maintenance. This research was funded in whole by the Austrian Science Fund (FWF) 10.55776/ZK74.

---

**Figure S1.**
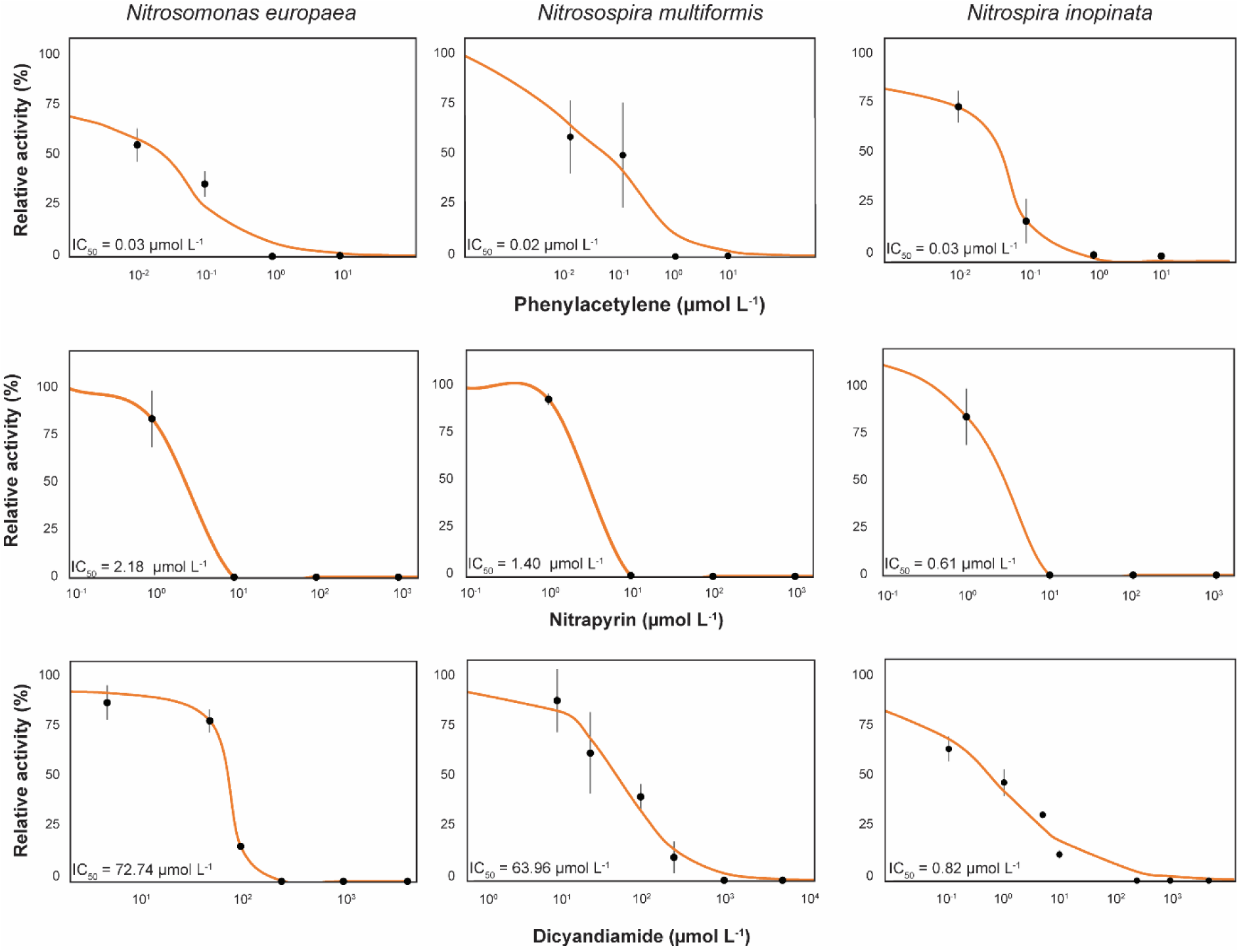
Constrained logistic regression models of growth inhibition when ammonium was provided as the sole substrate. Relative activity of *N. europaea* (left), *N. multiformis* (middle) and *N. inopinata* (right) cultures growing in the presence of different concentrations of phenylacetylene (top), nitrapyrin (middle) and dicyandiamide (bottom). IC_50_ values were modeled from dose-response curves using a logistic regression model (i.e., the Hill-Langmuir equation, Sebaugh, 2010). Error bars represent the standard deviation of three independent culture replicates.

**Figure S2.**
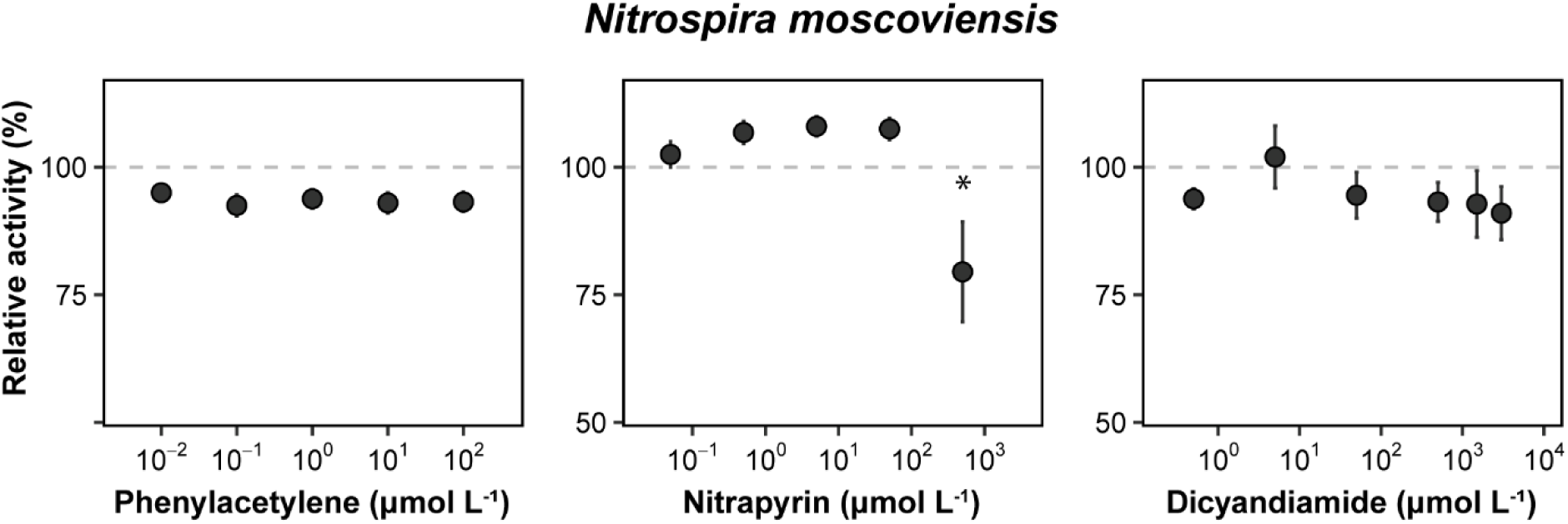
Sensitivity of *N. moscoviensis* to three common nitrification inhibitors. Relative respiratory activity of NOB *N. moscoviensis* culture, supplied with 1 mmol L^-1^ NO_2_^-^ as sole substrate, in response to injections of increasing concentrations of PA (left), NP (middle) and DCD (right). Error bars represent the standard deviation of three independent culture replicates. The asterisk indicates significant inhibition as determined by a paired t-test (p< 0.01).

**Table S1.** Experimental conditions for microrespirometry inhibition assays. Growth temperature, substrate concentration, and culture fold concentration used for microrespirometry-based inhibitor assays.

|  | Optimal growth temperature (°C) | When NH<sub>4</sub><sup>+</sup> or NO<sub>2</sub><sup>-</sup> was provided as the initial substrate |  | When N<sub>2</sub>H<sub>4</sub> was provided as the initial substrate |  | References |
| --- | --- | --- | --- | --- | --- | --- |
|  |  | Substrate concentration (mmol L<sup>-1</sup>) | Culture concentration (fold) | Substrate concentration (mmol L<sup>-1</sup>) | Culture concentration (fold) |  |
| *Nitrospira moscoviensis* | 37 | 1 | 5x | N/A | N/A | Nowka *et al.*, 2015 |
| *Nitrospira inopinata* | 37 | 0.3 | 15x | 0.3 | 47-95x | Kits *et al.*, 2017 |
| *Nitrosomonas europaea* Nm50 | 28 | 10-15 | 5x | 0.3 | 11-17x | Jung *et al.*, 2022 |
| *Nitrosospira multiformis* NI2 | 28 | 10 | 15x | 0.3 | 17-40x | Jiang and Bakken, 1999 |

**Table S2.**
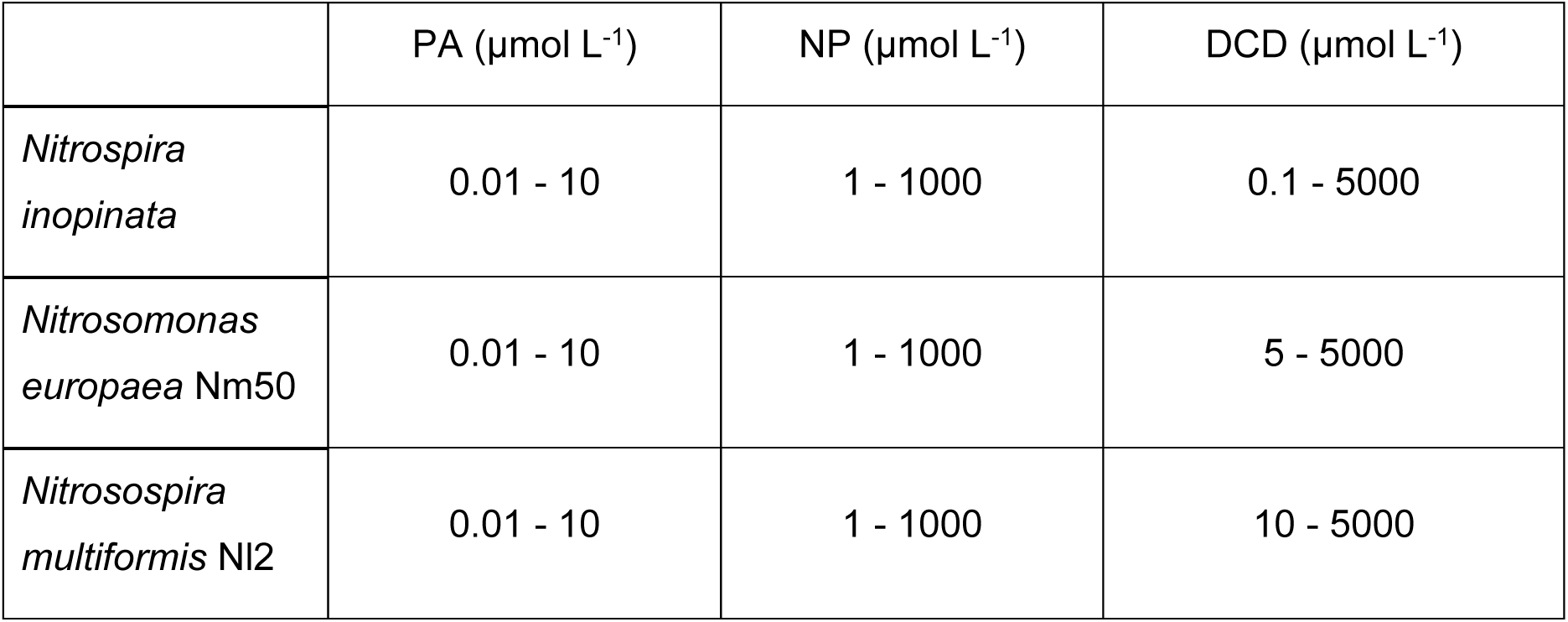
Inhibitor concentration range used for batch growth assays. PA-phenylacetylene; NP - nitrapyrin, DCD - dicyanamide. All assays were performed in the presence of 1 mmol L^-1^ NH_4_Cl provided as substrate, 0.1% v/v DMSO provided as solvent, in the dark, at the respective optimal growth temperatures listed in Table S1.

|  | PA (μmol L<sup>-1</sup>) | NP (μmol L<sup>-1</sup>) | DCD (μmol L<sup>-1</sup>) |
| --- | --- | --- | --- |
| *Nitrospira inopinata* | 0.01 - 10 | 1 - 1000 | 0.1 - 5000 |
| *Nitrosomonas europaea* Nm50 | 0.01 - 10 | 1 - 1000 | 5 - 5000 |
| *Nitrosospira multiformis* NI2 | 0.01 - 10 | 1 - 1000 | 10 - 5000 |

